# Transendothelial migration induces differential migration dynamics of leukocytes in tissue matrix

**DOI:** 10.1101/2021.09.02.458715

**Authors:** Abraham C.I. van Steen, Lanette Kempers, Rouven Schoppmeyer, Max Blokker, David J. Beebe, Martijn A. Nolte, Jaap D. van Buul

**Author notes:** **Correspondence should be addressed to:** Jaap D. van Buul, Molecular Cell Biology Lab, Dept. Molecular Hematology, Sanquin Research and Landsteiner Laboratory, Academic Medical Center at the University of Amsterdam, Plesmanlaan 125, 1066CX Amsterdam, the Netherlands. These authors contributed equally (Abraham C.I. van Steen, Lanette Kempers).

## Abstract

Leukocyte extravasation into inflamed tissue is a complex process that is difficult to capture as a whole *in vitro*. We employed a blood-vessel-on-a-chip model in which endothelial cells were cultured in a tube-like lumen in a collagen-1 matrix. The vessels are leak-tight, creating a barrier for molecules and leukocytes. Addition of inflammatory cytokine TNF-α caused vasoconstriction, actin remodelling and upregulation of ICAM-1. Introducing leukocytes into the vessels allowed real-time visualisation of all different steps of the leukocyte transmigration cascade including migration into the extracellular matrix. Individual cell tracking over time distinguished striking differences in migratory behaviour between T-cells and neutrophils. Neutrophils cross the endothelial layer more efficiently than T-cells, but upon entering the matrix, neutrophils display high speed but low persistence, whereas T-cells migrate with low speed and rather linear migration. In conclusion, 3D imaging in real-time of leukocyte extravasation in a vessel-on-a-chip enables detailed qualitative and quantitative analysis of different stages of the full leukocyte extravasation process in a single assay.

**Summary:** A functional hydrogel-based blood-vessel-on-a-chip model is used to study the complete leukocyte transendothelial migration process in real time. T-lymphocytes and neutrophils exhibit distinct migration dynamics in the extravascular matrix after transendothelial migration, which can be altered using a chemotactic gradient.

## Introduction

Leukocyte transendothelial migration (TEM) forms the basis of immune surveillance and pathogen clearance, and hence plays a pivotal role in many (patho)physiological processes. This process is highly regulated and strongly differs between the various leukocyte subsets and tissues involved. Leukocyte extravasation mainly occurs in postcapillary venules, where adherens junctions allow for transient opening of the barrier to allow leukocytes to pass.

Importantly, most venules only allow leukocyte TEM when the surrounding tissue is inflamed, thereby locally exposed to pro-inflammatory cytokines. One of those is tumour Necrosis Factor-α (TNF-α), which is produced by a variety of cells under inflammatory conditions (Heller and Krönke, 1994). Once leukocytes have crossed the endothelial barrier, they continue migrating into the underlying matrix to fight the invading pathogens (Woodfin, Voisin and Nourshargh, 2010). Tissue penetration is an important aspect of the extravasation process as this determines if a pathogen will be successfully cleared or not (Yamada and Sixt, 2019). It is therefore essential to understand how immune cells migrate through the extracellular matrix once they have crossed the endothelium. To date, the full extravasation process, including intraluminal rolling, crawling, diapedesis and the extracellular 3D matrix migration, can only be studied using *in vivo* models as no proper *in vitro* models are available. As *in vivo* models have their limitations, it would be more than desirable to have an *in vitro* system that allow to study the full process in real-time.

Research regarding the TEM process on a cellular level is mainly done using 2D *in vitro* models, generally based on endothelial cell monolayers cultured on flat stiff surfaces, such as coverslips (Muller and Luscinskas, 2008). Substrate stiffness has been shown to affect cell-cell and cell-matrix interactions of endothelial cells, as well as leukocyte–endothelial cell interactions, and subsequently TEM (Huynh *et al.*, 2011; Stroka and Aranda-Espinoza, 2011). Another limitation is that leukocytes encounter this stiff impermeable surface after traversing the endothelial barrier, making it impossible to study leukocyte detachment from the vessel and entry into the tissue. 3D systems, such as Transwell assays and Boyden chamber assays, allow leukocytes to advance beyond the endothelial cell monolayer, however these systems are unsuitable for microscopy-based imaging and are also based on stiff substrates (Muller and Luscinskas, 2008). To achieve the next step in TEM research, a more intricate model is required that allows imaging of the entire process from the luminal side of the endothelium into a physiological matrix substrate.

Organ-on-a-chip (OOAC) models use microfluidics-based approaches to create biomimetic systems emulating physiological organ function. Blood vessels are well suited for OOAC development, as their structure is relatively simple compared to entire organs (Virumbrales-Muñoz *et al.*, 2020). Knowledge on vascularisation of *in vitro* systems obtained from Blood-Vessel-on-a-Chip (BVOAC) models could also be applied to organoids to overcome their current size and function limitations. BVOAC systems typically consist of a tubular EC monolayer generated in a stiff glass/plastic substrate (Farahat *et al.*, 2012; Zervantonakis *et al.*, 2012; Zheng *et al.*, 2012) or hydrogel (Wong *et al.*, 2012; Kim *et al.*, 2016; Sobrino *et al.*, 2016). A perfusable and hydrogel-based BVOAC system overcomes the aberrations associated with stiff substrates and in addition allows leukocytes to extravasate into the matrix surrounding the vessel.

A system that meets these requirements is the LumeNext system, which is also highly suited for imaging analysis (Jiménez-Torres *et al.*, 2016). Apart from building a 3D blood vessel in a physiological matrix, this device allowed us to track the subsequent migration of primary human neutrophils as well as T-cells beyond the diapedesis stage into the matrix in real-time using confocal microscopy. We discovered that neutrophils and T-Lymphocytes use markedly different migration modes to enter the tissue. Moreover, by applying a chemotactic gradient of complement component 5a (C5a) into the hydrogel matrix, we found that C5a did not increase the number of neutrophils that crossed the endothelium, but rather promoted migration towards C5a by adjusting directionality of cell migration, while reducing migration speed. These findings demonstrate that the combination of cutting-edge BVOAC systems with advanced microscopy offers valuable new insights into how different leukocyte subsets respond to chemokines and cross the vascular barrier and enter the tissue.

## Results

### Characterization of endothelium-lined blood vessels

To study leukocyte TEM in a reproducible BVOAC device and monitor this in real time, we used the LumeNext device (Figure 1A) (Jiménez-Torres *et al.*, 2016). Endothelial cells were seeded in the lumen inside a collagen-1 matrix using a head-over-head incubator, allowing the formation of a 3D vessel (Figure 1B). Staining of VE-cadherin, F-actin and nuclei showed the formation of a confluent endothelial monolayer (Figure 1C), as well as a vascular lumen (Figure 1D). Detailed imaging showed a confluent endothelial monolayer with linear junctions, marked by VE-cadherin and F-actin, representative for a functional barrier (Figure 1E) (Ando *et al.*, 2013).

**Figure 1.**
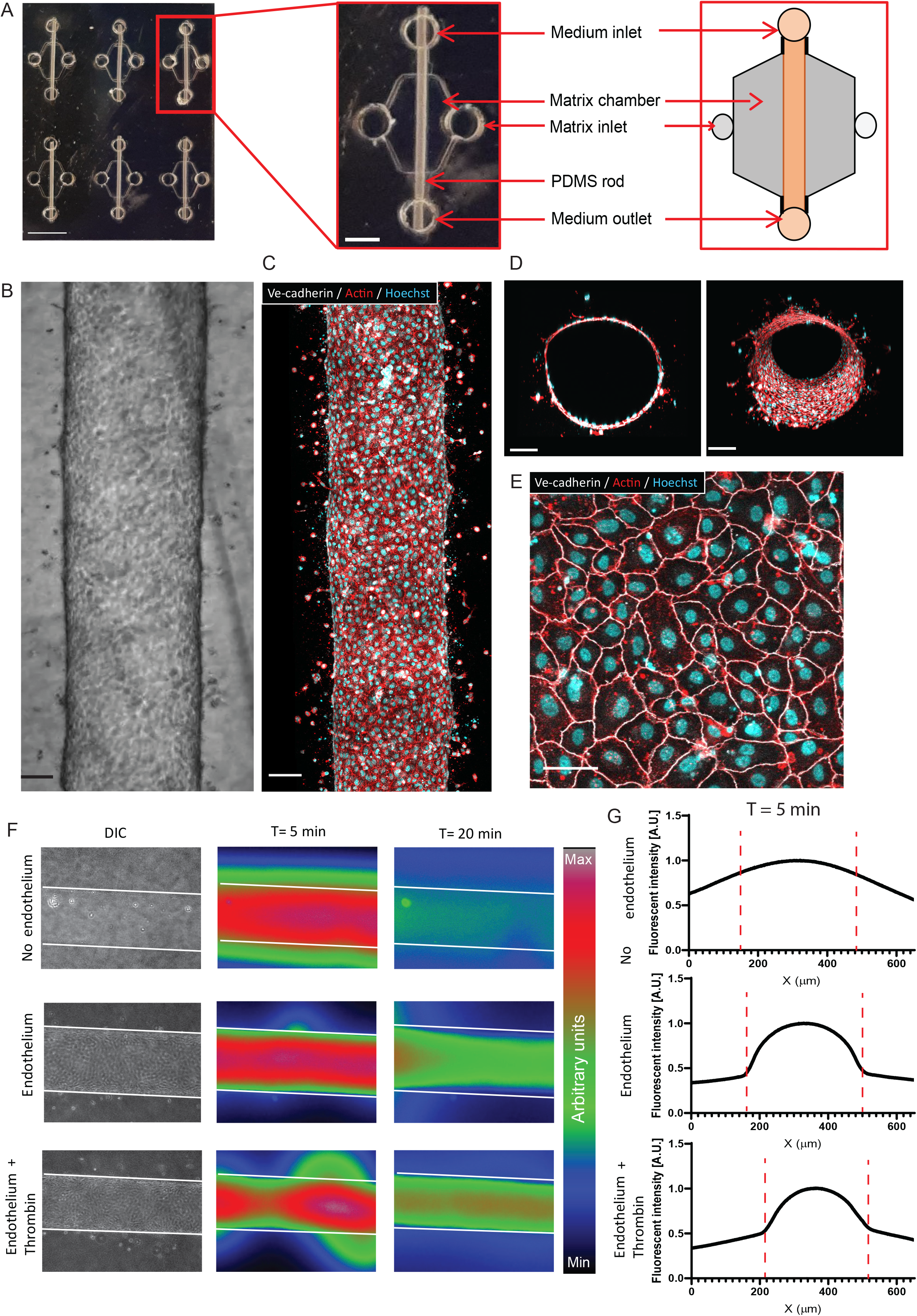
Making and characterization of the BVOAC. Overview of the device (A) containing six chambers to make a BVOAC, zoom-in on one BVOAC and schematic overview of one vessel place. DIC (B) and confocal (C) image of a lumen lined with endothelial cells (D) and orthogonal view showing the open lumen of the BVOAC and the Imaris surface rendering. Zoom in on the BVOAC in D showing the endothelial monolayer (E). Representative images of leakage of 70 kDa dextran in the BVOAC (F) and quantification of the leakage 5 min after injection (G). Quantification was done by measuring the average fluorescent intensity over the Y axis of the image which is then normalised to the maximum signal measured. The red dotted lines represent the vessel outlines. Scalebar = 3 mm + 1 mm (A), 100 μm (B, C, D), 50 μm (E).

As the barrier function of endothelial monolayers is an essential function of blood vessels, we measured the barrier function in the BVOAC model by perfusing the lumen with fluorescently-labelled 70 kDa dextran, comparable size to albumin, an abundant plasma protein. In a non-vascularized lumen (i.e., no endothelial lining), dextran rapidly diffused into the matrix, whereas in endothelialised lumen, dextran was contained for more than 20 min without leakage. Addition of thrombin, a well-known vascular permeability factor (Van Nieuw Amerongen *et al.*, 1998) to the lumen temporally induced endothelial permeability that recovered over time (Figure 1F-G). This shows that the vessels are functionally lined with endothelial cells and provide a proper barrier function.

### BVOAC under inflammatory conditions

Upon inflammation the endothelium upregulates crucial adhesion molecules and chemokines required for leukocyte extravasation. Therefore, we incubated the vessels overnight with the inflammatory mediator TNF-α to activate the endothelial cells (Figure 2A-B). Interestingly, TNF-α did not alter the number of endothelial cells in the BVOAC (Figure 2C) but did lead to a smaller vessel diameter 24 hours after addition of TNF-α (Figure 2D). TNF-α treatment resulted in the induction of actin stress fibres through the cell body and the upregulation of ICAM-1 (Figure 2E). In untreated cells, F-actin was predominantly present at the cell junctions where they colocalised with VE-cadherin. In TNF-α treated cells, cytosolic actin stress fibres are observed in addition to junctional actin stress fibres (Figure 2F). In addition, the endothelial cells produced a layer of Collagen IV around the vessel, making up a basement membrane (Figure 2G). In the orthogonal view, it can be observed that the Collagen IV is present at the baso-lateral side of the endothelium (Figure 2H). The reduced vessel size, upregulation of ICAM-1, increased stress fibre formation and presence of Collagen IV around the vessel illustrate that the endothelial cells were in an inflamed state after TNF-α treatment.

**Figure 2.**
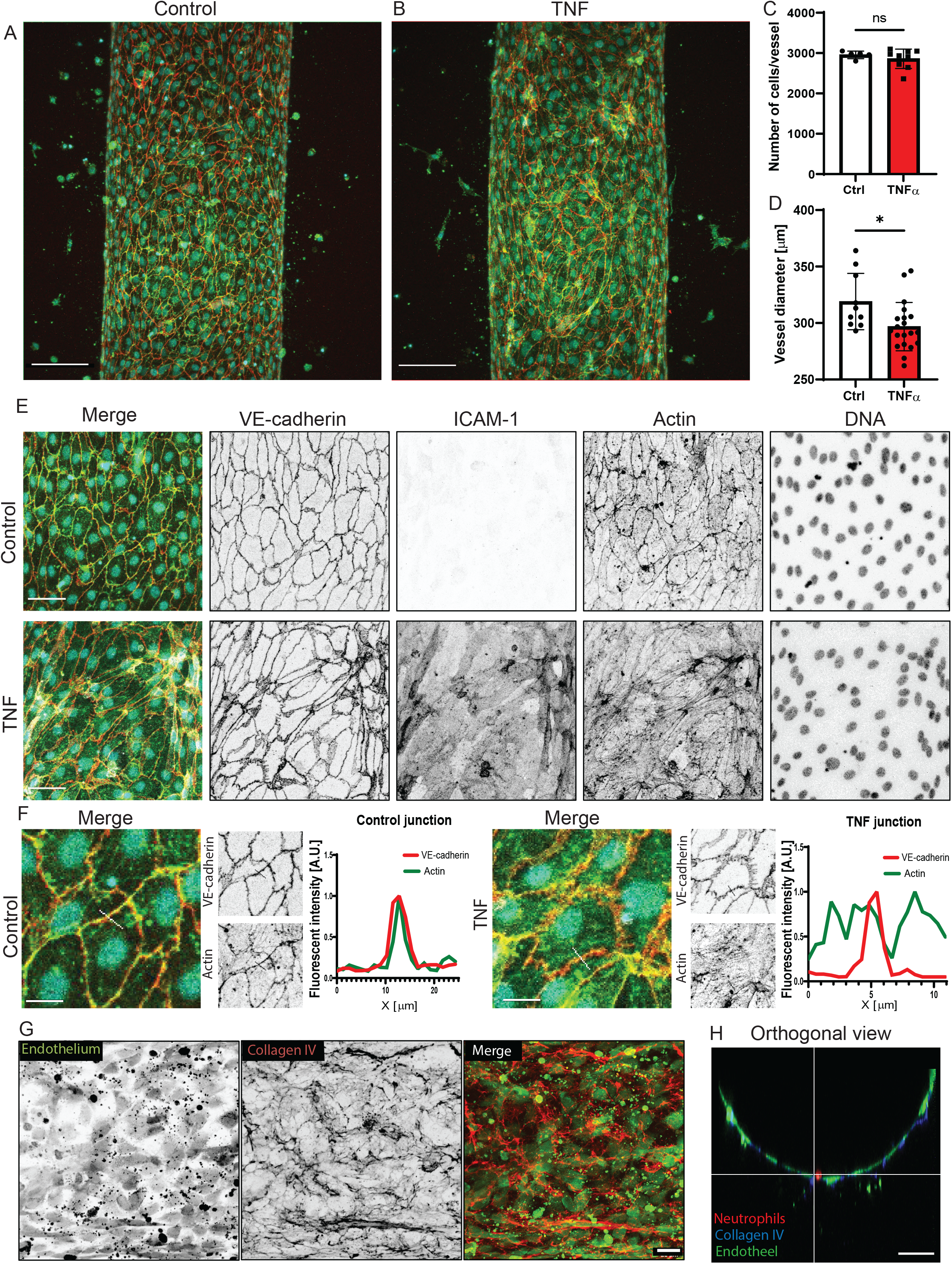
TNF-α in the vessel induces inflammation. Overview of a control (A) or TNF-α treated vessels (B). Quantification of the number of cells (C) and vessels diameter (D) in TNF-α treated vessels compared to control vessels. Representative images of control and inflamed endothelial cells (E) and junctions with line plots showing the localization of VE-cadherin and actin in these junctions (F). Representative image of the presence of Collagen IV around the vessels (G). Orthogonal view of a vessel with collagen IV staining around (H). Scalebar = 100 μm (A, B), 50 μm (E, H), 25 μm (F, G). Data quantified from 5 (Ctrl), 9 (TNF-α) BVOAC for the number of ECs per BVOAC, diameter 10 (Ctrl) and 20 (TNF-α) BVOAC, staining’s and line plots 3 BVOAC per condition.

### 3D analysis of leukocyte transmigration in vessels

Next, we investigated TEM of neutrophils in the BVOAC model. Neutrophils were injected into uninflamed and inflamed vessels and incubated for 2.5 hours. The devices were then washed to remove non-adherent neutrophils, fixed, and imaged using confocal microscopy. We found that neutrophils not only crossed the endothelial monolayer but also continued migration into the underlying collagen matrix (Figure 3A). Imaris software was used to generate a 3D rendering of the BVOAC, which revealed neutrophils mostly transmigrated at the bottom of the vessel (Figure 3B) due to gravity. When allowing transmigration during continuous turning of the device, thereby reducing the effect of local gravity, we found that neutrophils left the vessels at all sides (Figure S1). This rendering was adapted for analysis by manually creating a surface at the position of the vessel (Figure 3C) and creating spots at the centre of mass for neutrophils (Figure 3D), based on automatic detection and manual curation. Following this analysis step, we were able to calculate the distance between the surface and the individual spots who were then colour coded, with warm colours representing longest distance from the endothelium into the matrix (Figure 3E). This analysis allowed us to distinguish between the different steps of TEM, namely adhesion, diapedesis, and penetration into the ECM (Figure 3G). To demonstrate the three different steps of the TEM cascade, we imaged in more detail the wall of the vessel and were able to observe the full TEM process, i.e., rolling, adhesion, and diapedesis (Figure 3H, Supplemental Video 1-2). In the video 2 neutrophils transmigrate at the same spot, shortly after each other, indicating the possible presence of a transmigration hotspot (Grönloh, Arts and van Buul, 2021). Quantification showed that the distance neutrophils travelled from the vessels into the matrix ranging from −18 μm (in the lumen) to 286 μm into the matrix (Figure 3F). Neutrophils with a distance of ≤ 8 μm from the vessel were still adhering to the endothelium. From these data, we conclude that this new analysis tool can track individual neutrophils at all stages of TEM, comparable with *in vivo* analysis.

**Figure 3.**
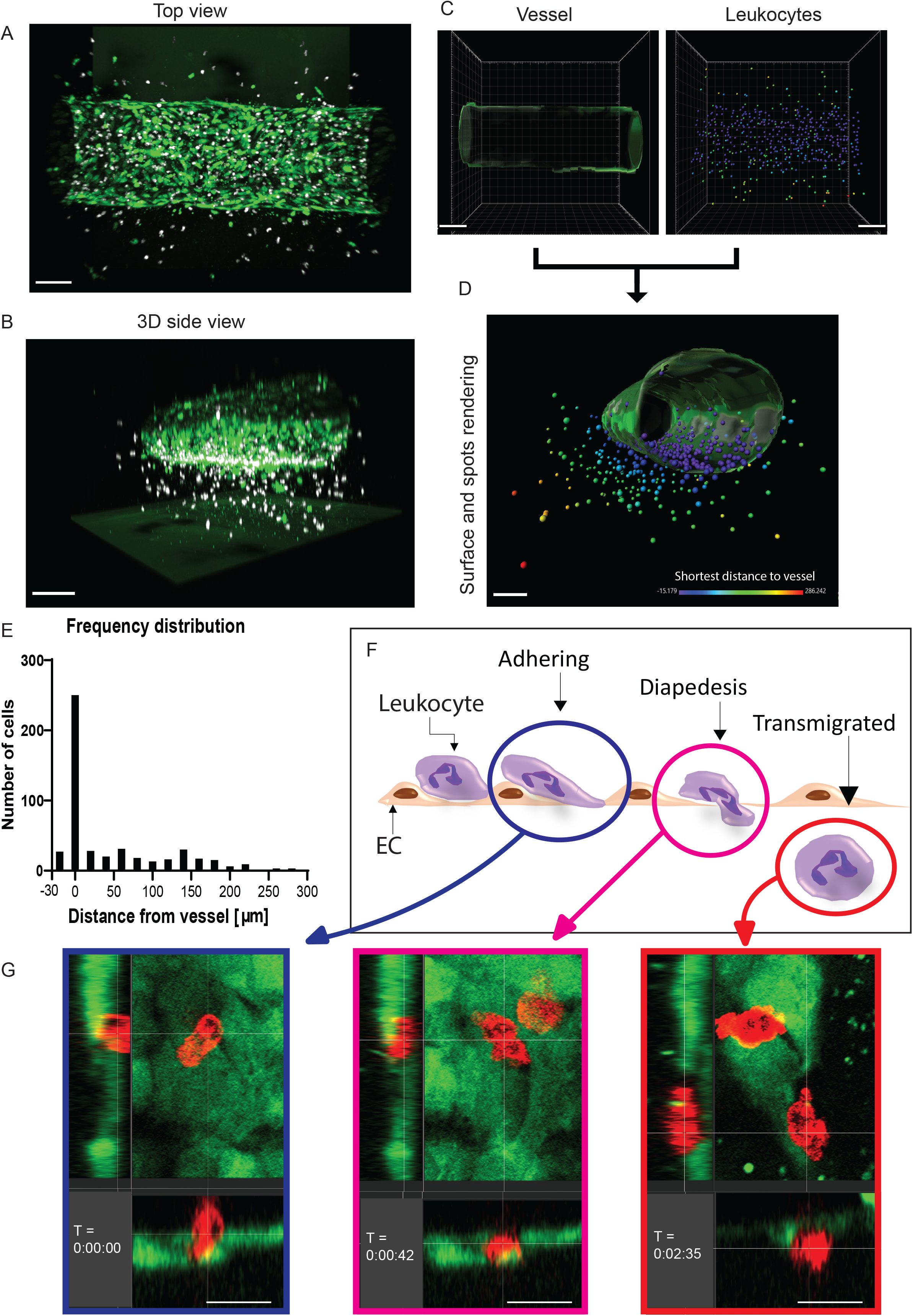
Vessel and leukocyte analysis in Imaris. Microscopy image depicting the BVOAC (green) with the neutrophils (white) from top (A) and side view (B). Surface of the BVOAC (C) and the leukocytes (D). Overview of the BVOAC surface with colour coded spots based on the distance from surface allowing calculations and analysis (E). Distribution of the distance between the leukocytes and the vessel displayed in E (F). Schematic overview of the different transmigration steps: adhering to the endothelial cells on the inside of the lumen (Blue circle), diapedeses (Pink circle) and transmigrated cells (Red circle) (G). Orthogonal view of stills taken from detailed high-speed imaging of neutrophil TEM in the vessel over time (H). Scalebar = 150 μm (B-D), 100μm (A,E), 15 μm (H).

### BVOAC allows chemokinesis-mediated TEM into the tissue

To quantify the number of neutrophils that left the vessel lumen and entered the matrix, we used a 3D rendering and tracking approach to study chemokine-induced neutrophil TEM under inflammatory conditions. This 3D rendering allowed us to observe a difference in number of transmigration events that was not detectable using a 2D analysis (Figure 4A-B). Using this approach, we accurately discriminated between adherent and transmigrated neutrophils (Figure 4A), revealing a strong increase in both leukocyte TEM states in TNF-α-stimulated vessels compared to control vessels (Figure 4C). Interestingly, neutrophils that crossed TNF-α-stimulated endothelium migrated further away from the vessel than neutrophils that crossed untreated endothelium, indicating that the inflamed endothelium may stimulate neutrophil migration (Figure 4D).

**Figure 4.**
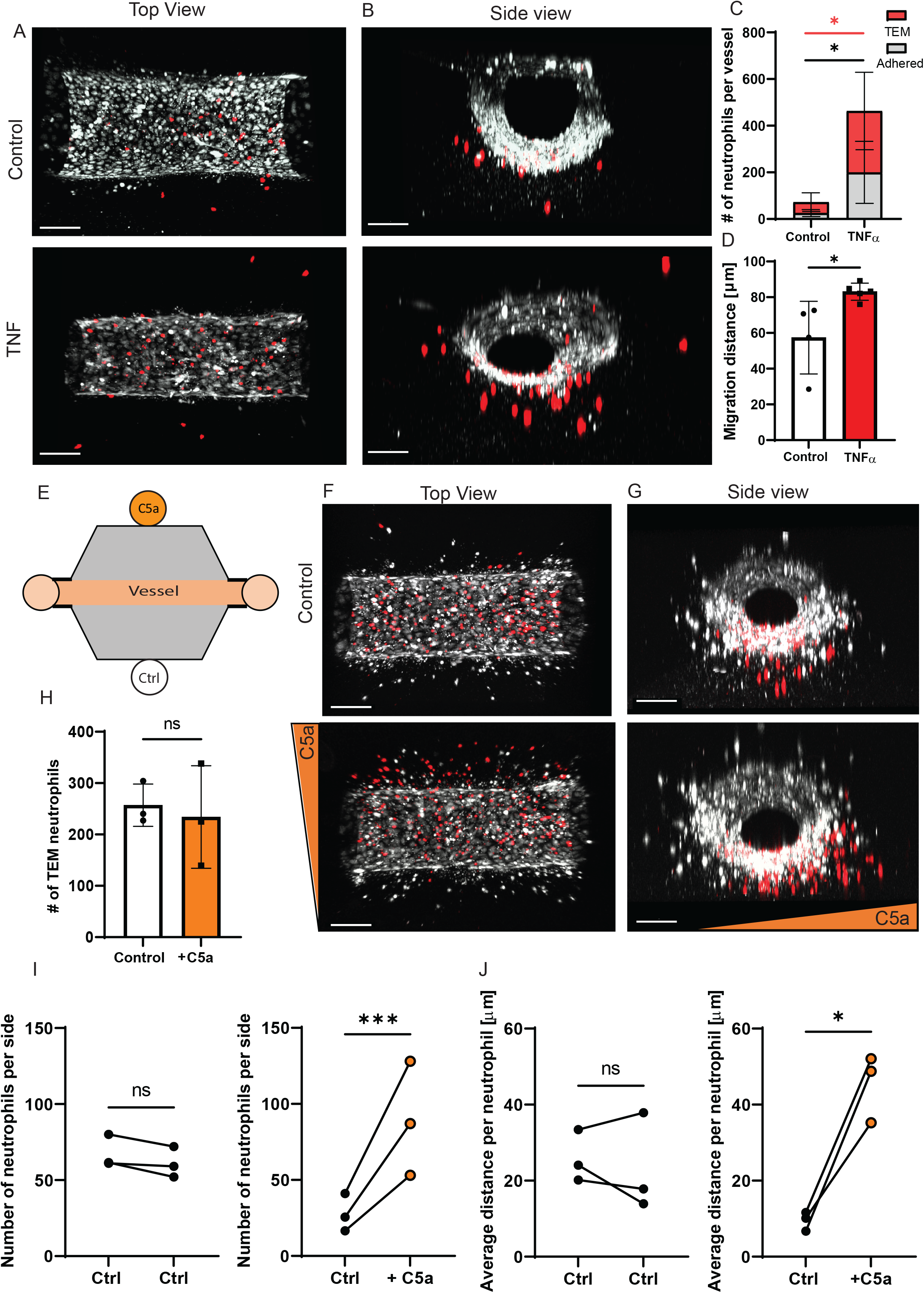
Neutrophil transmigration in a 3D environment. Representative top view (A) and side view (B) images of neutrophil transmigration in control and TNF-α treated BVOAC. Quantification of the number of neutrophils per field of view (C) and average transmigration distance after exiting the BVOAC (D). Diagram of the experiment with one-sided addition of C5a (E). Representative top view (F) and side view (G) images of neutrophil transmigration in TNF-α treated BVOAC with C5a or PBs on one side. Quantification of the total number of transmigrated neutrophils per field of view (H), # of neutrophils per side of PBS and C5a treated BVOAC (I) and average migration distance between left and right of PBS or C5a treated BVOAC (J). Scalebar = 150 μm. Data quantified from 4 BVOAC per condition for neutrophil TEM and 3 BVOAC per condition for C5a TEM.

To test if neutrophils respond to a chemotactic gradient, we injected C5a, a chemoattractant, into one of the matrix inlets (see schematic Figure 4E). Addition of C5a did not alter the number of neutrophils that crossed inflamed endothelium (Figure 4H) but changed the migration direction of neutrophils towards C5a significantly (Figure 4F-G, I). Interestingly, whereas the migration distance that neutrophils travelled was not different between the two sides under control conditions (Figure 4J-left), in the presence of C5a on one side the migration distance towards this side was significantly increased (Figure 4J-right). In summary, these data demonstrate that luminal application of TNF-α in this BVOAC model strongly enhances neutrophil extravasation and entry into the surrounding matrix, which can be directionally steered by applying a chemotactic gradient inside the matrix.

### T-cell transmigration through a HUVEC lined vessel and neutrophils transmigration in pancreatic and lung microvascular vessels

To examine the versatility of this system and determine whether this system is also suitable for analysing T-cell TEM, we injected purified T-cells into the lumen of both inflamed and control vessels. After 2.5 hours, vessels were flushed, fixed, and imaged (Figure 5A-B). We found that the number of adhered and transmigrated T-cells was 5 times increased in inflamed vessels compared to control conditions (Figure 5C). In addition, T-cells migrated further into the underlying matrix when the vessels were inflamed (Figure 5D), in line with the finding that neutrophils also extended their migration track when crossing inflamed endothelium. However, whereas neutrophils travelled from the vessel into the matrix for on average 83 μm, T-cells only travelled on average 44 μm, indicating that the intrinsic migration speed of neutrophils through the 3D matrix is 89% higher compared to T-cells.

**Figure 5.**
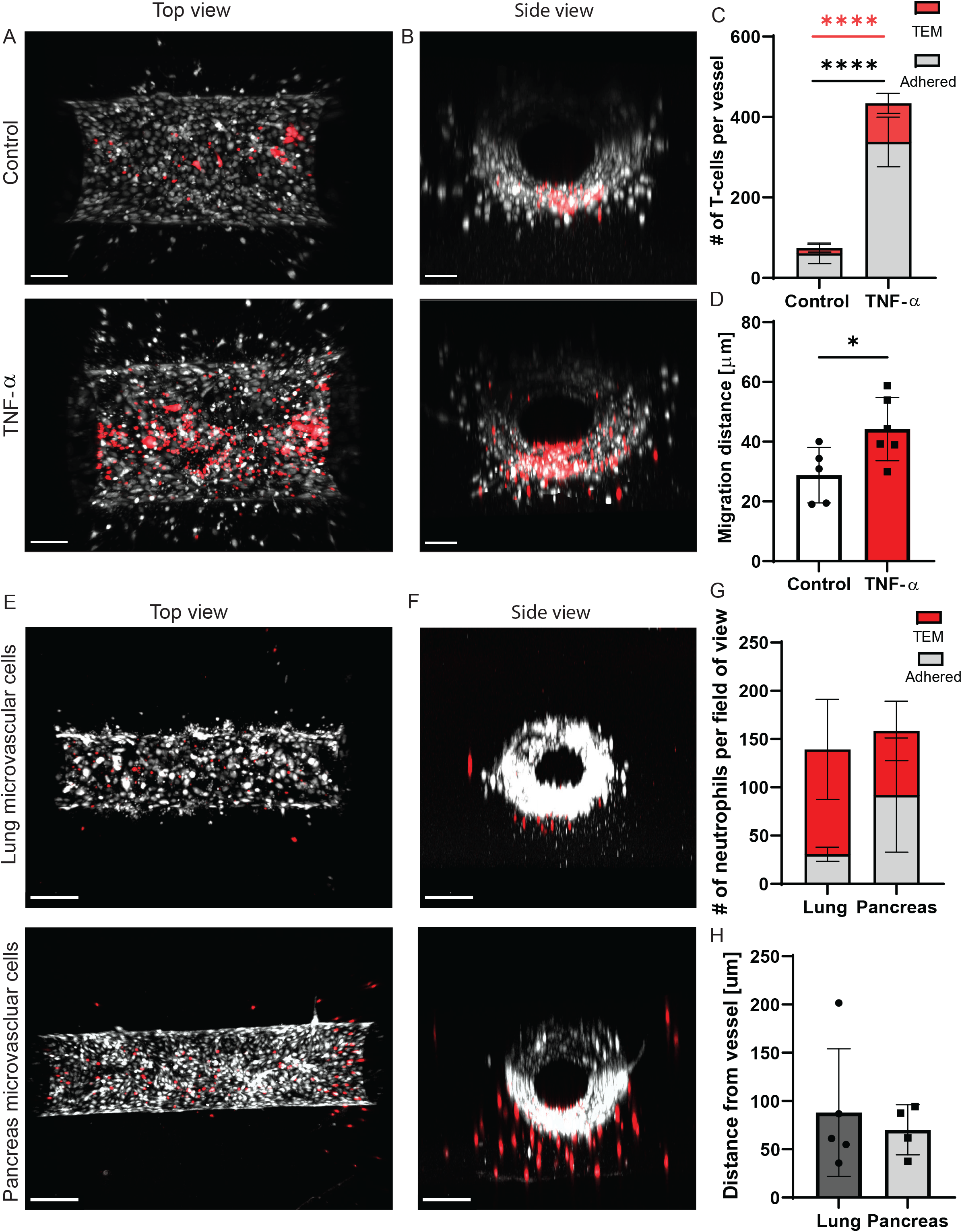
T-cell transmigration in a 3D-environment. Representative top view (A) and side view (B) images of T-cell transmigration in control and TNF-α treated BVOAC. Quantification of number of transmigrated cells (C) and cumulative transmigration distance (D). Representative top view (E) and side view (F) images of neutrophil transmigration in vessels lined with lung or pancreatic endothelial cells. Quantification of number of adhered and transmigrated cells (G) and migration distance of the transmigrated cells (H). Scalebar = 50 μm. Data quantified from 4 (pancreas), 5 (Ctrl, lung vessels) and 6 (TNF-α) vessels per condition.

In addition to using different leukocyte subsets, we analysed if different endothelial cells may be used to generate the vessel. For this, we used human microvascular endothelial cells from both the lungs and pancreas. Both cell types formed a nice vessel (Figure 5E-F). Next, neutrophils were added to the vessels of both endothelial cell types, similar to previous experiments. Neutrophils transmigrated across both endothelial cell types and migrated into the matrix (Figure 5G). Although the total number of cells in the field of view did not differ between lung and pancreas vessels (138 and 158 respectively), transmigration percentage was increases in the lung vessels (78% and 41% respectively). The distance travelled by neutrophils from the vessel into the matrix was similar in both cell types (Figure 5H). These data indicate that the BVOAC device is very versatile, both for different leukocyte and endothelial cell types, and can be adjusted to the research question that needs to be answered.

### Live cell imaging reveals differences in leukocyte migration

We next investigated the source of the difference in migration distance between T-cells and neutrophils. Although 3D confocal imaging analysis offers a more accurate detection of transmigration events compared to 2D widefield imaging, it does not allow us to distinguish between actual migration speed and relative track distance over time. Using high-speed imaging, we generated Z-stacks of 200 μm with a time interval of 10 seconds, giving us sufficient spatiotemporal resolution for semi-automatic tracking of individual leukocytes through the collagen matrix in time. Automatic tracking was performed on all leukocytes in the field of view, after which tracks were filtered to exclude neutrophils exiting or entering the field of view. For analysis requiring complete leukocyte tracks, such as displacement or total track length, we selected 10 tracks per experiment which were manually checked to assure quality of the data.

Using this approach, we were able to track migrating neutrophils or T-cells in time in 3 dimensions (Figure 6A-B and Supplemental Video 3-4). We found that the migration speed of T-cells inside the matrix after crossing the inflamed endothelium is significantly slower on average than that of neutrophils (Figure 6C; 0.10 μm/sec ± 0.03 and 0.20 μm/sec ± 0.02, respectively). Based on these measurements neutrophil migration speed is 100% higher compared to T-cells, which is different from the 89% larger migration distance observed in the endpoint experiment. To assess whether the speed of both leukocyte subsets was consistent over time from the initial start until the end of the recordings, we compared the initial and final migration speed, and found that both neutrophils and T-cells maintain a constant speed over time (Figure 6D-E).

**Figure 6.**
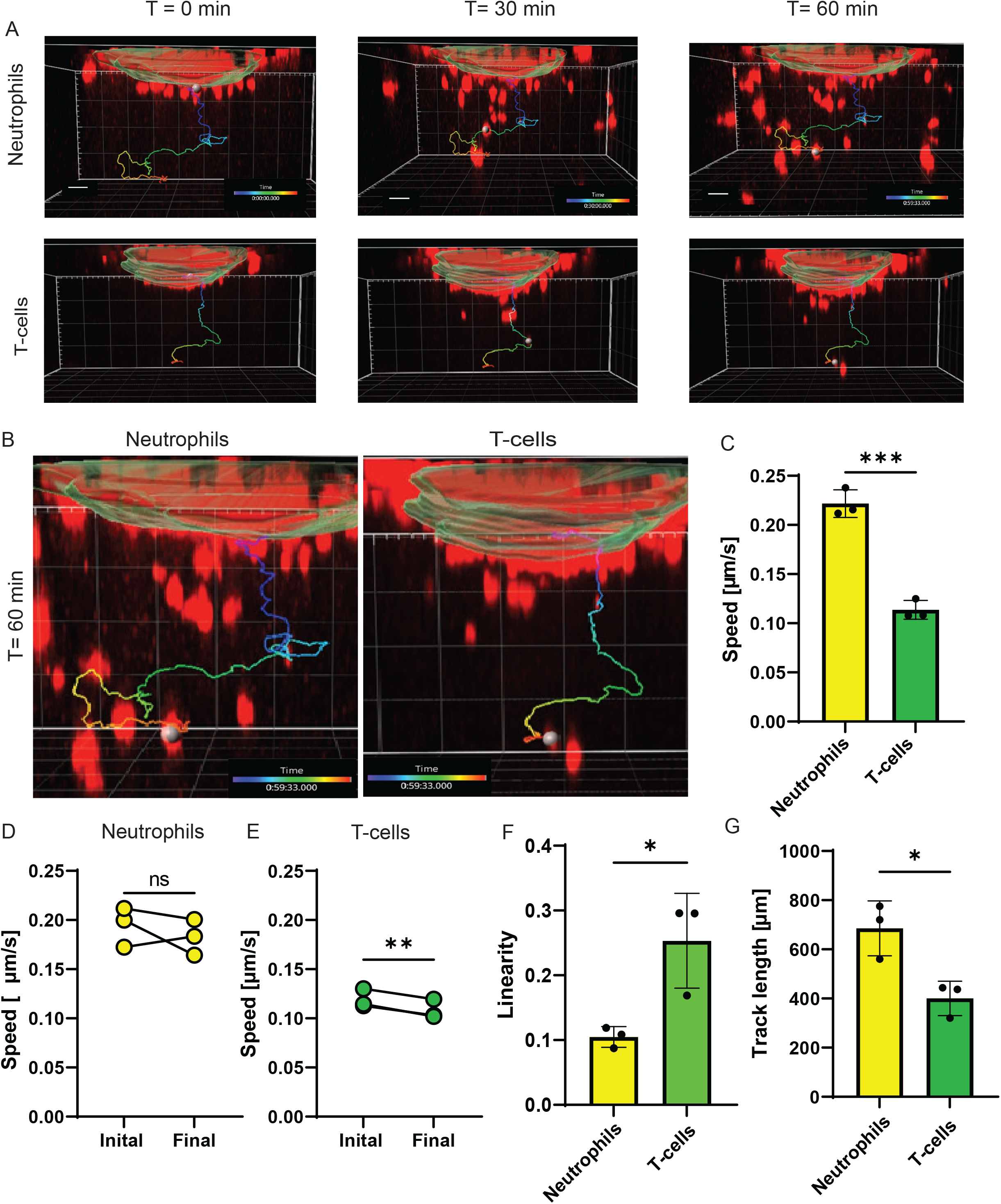
Live transmigration of neutrophils and T-cells. (A) Stills from a video in which neutrophils and T-cells migrate out of the BVOAC. Per condition one track representing the general migration manner of that leukocyte is highlighted. (B) Zoom-in on the track at T=60 of both neutrophils and T-cells. Quantification of the average track speed (C), initial and final migration speed of neutrophils (D) and T-cells (E), linearity (F), track length (G) for both neutrophils and T-cells. Scalebar = 50 μm. Data quantified from 10 cells per BVOAC and 3 BVOAC per condition.

Next, we analysed the linearity of the migration tracks, which is defined as the ratio of the displacement over the total distance travelled by that cell (Boissonnas *et al.*, 2007). This ratio has a maximum of 1, which occurs when the cells travelled in a straight line from the start position to the end position. We found that T-cells migrate in a more linear manner compared to neutrophils (Figure 6F; 0.25 ± 0.07 and 0.10 ± 0.02, respectively). Accordingly, the total distance travelled within the same time frame was higher for neutrophils than for T-cells (Figure 6G; 948 μm vs. 400 μm, respectively). These results indicated that neutrophils exerted a more intrinsic exploratory behaviour than T-cells in our system. Therefore, we conclude that neutrophils migrate faster but wander around more through the collagen, whereas T-cells migrate slower, but in a more directed fashion.

### C5a gradient increases neutrophil speed in the first half hour

We examined to what extent the exploratory behaviour of neutrophils can be regulated by a chemotactic gradient such as C5a. 3D live imaging of neutrophil migration in the presence of a C5a gradient showed that most neutrophils migrated towards the chemoattractant (Figure 7A-B, Supplemental Video 5), confirming our previous results. Interestingly, when analysing the direction of migration over time, we observed that the C5a-driven migration pattern occurred predominantly in the first 15 min of the experiment. After this timepoint, C5a-driven migration directionality was strongly diminished and neutrophil migration directionality appeared to be random, as if no chemoattractant was present (Figure 7C-D). We also calculated the speed of the neutrophils that were exposed to C5a and found that over the course of the experiment, the speed was reduced (Figure 7E). We found that the migration linearity was significantly increased in the presence of C5a compared to control condition (Figure 7F; 0.17 ± 0.04 vs. 0.10 ± 0.02, respectively). The decreased migration speed and increased directionality together lead to a total migration distance that was not changed in the presence of C5a compared to control (Figure 7G). These data indicate that the chemotactic gradient in the matrix disappeared in time, with the consequence that neutrophils change their migratory behaviour from linear to random migration pattern.

**Figure 7.**
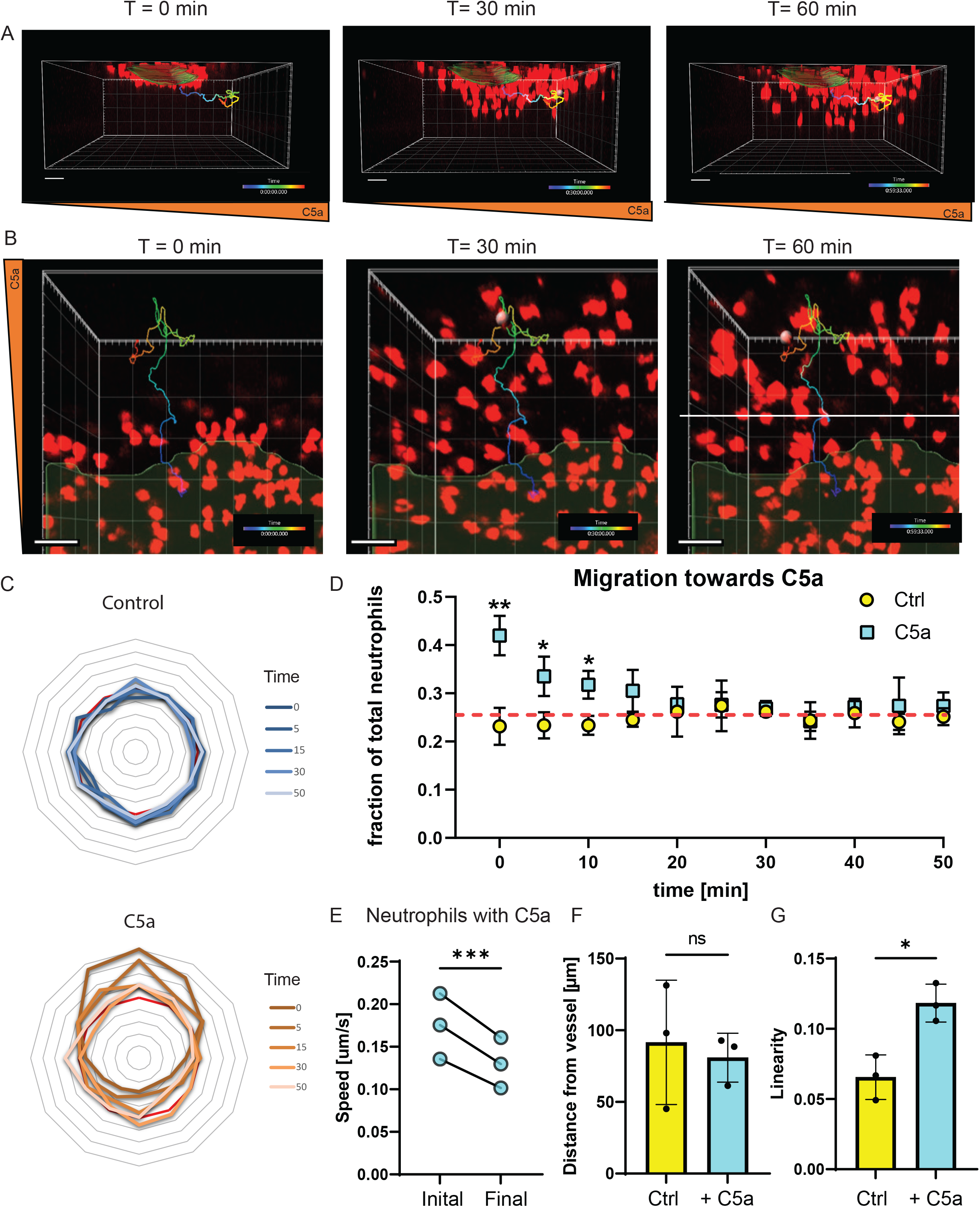
Live transmigration of neutrophils and migration dynamics with C5a. Stills from a movie showing the migration of neutrophils towards one side with a side (A) and top (B) view. Quantification of the direction of the neutrophils in control and the C5a condition over time (C, D). This windrose plot shows the distribution of leukocyte migration directions at distinct times as shown in the legend. Each concentric axe indicates 1.7% of the total number of neutrophil movements at the indicated time. The red line represents the situation where leukocyte migration in every direction is equal, and directionality is not observed. Quantification of the initial and final speed of neutrophils in the presence of C5a (E), endpoint distance from the vessel (F) and the ratio between the end point distance from the vessel and the total length travelled, also called linearity (G). Scalebar = 50 μm. Data quantified from 10 cells per BVOAC and 3 BVOAC per condition.

In conclusion, this BVOAC device enables us to accurately visualize and analyse TEM of primary human leukocytes in 3D over time, which can be modulated by applying a chemoattractant gradient. As such, this novel platform holds great promise for future studies unravelling the complex cellular and molecular mechanisms that underlie leukocyte TEM.

## Discussion

Leukocyte extravasation through the vessel wall is mostly studied using classical *in vitro* 2D models that allow for analysis of the molecular details of this process. However, in these models, endothelial cells are typically cultured on artificial stiff substrates, which alters endothelial behaviour and responses. Moreover, these models preclude analysis of the final part of the leukocyte extravasation; penetration of the surrounding matrix. Thus, there is an urgent need for a proper *in vitro* model that mimics the *in vivo* conditions and allows for studying of molecular details of this process. We demonstrate that a hydrogel-based BVOAC model perfectly meets these demands and allows studying the full extravasation process at the single cell level in 3D over time.

In the human body, inflammation typically leads to leads to vasoconstriction by contracting smooth muscle cells around the vessel wall (Pleiner *et al.*, 2003; Lim and Park, 2014). Leukocyte extravasation mostly occurs in inflamed post-capillary venules, where the blood vessels only consist of a single layer of endothelial cells (Baluk *et al.*, 1998). Because inflammatory mediators such as TNF-α induce strong F-actin stress fibres, they are expected to induce cellular tension and potential contraction on an endothelial monolayer (Wójciak-Stothard *et al.*, 1998). In 2D monolayers, TNF-α stimulation leads to the formation of intracellular gaps, an observation which is supported by an increase in permeability. However, using the BVOAC model, we confirmed that TNF-α increased the number of F-actin stress fibres, and the vessel lumen diameter was reduced, instead of a loss of contact between individual endothelial cells and the formation of intracellular gaps. In addition, in the human body, vessels are surrounded by a collagen IV layer (Yurchenco, 2011; Xu and Shi, 2014). In our system, this collagen IV layer was also present, mimicking *in vivo*-like conditions. This indicated that the BVOAC much better represents the physiological situation than any 2D cell culture model used to study inflammation-related events.

In addition, when culturing endothelial cells on glass or plastics, F-actin stress fibres are prominently present through the cell body, even without inflammatory stimuli. In human arteries, these fibres are also prominently present, however, in veins and post-capillary venules such fibres are lacking (Van Geemen *et al.*, 2014). Under control conditions, the BVOAC model represents the *in vivo* non-inflamed condition, as we observed no endothelial F-actin stress fibres. Only upon stimulation with TNF-α stress fibres appear in the cell body. Therefore, we conclude that our *in vitro* generated vessels accurately represent both resting and inflamed states of blood vessels *in vivo*, and thus are a major improvement compared to conventional inflammatory 2D models.

Another point where the BVOAC system shows its potential as a model is importance of real-time analysis of migrating leukocytes in 3D. When comparing end point measurements with real-time TEM data, we found that our analysis of fixed samples leads to a large underestimation of the migration distance, particularly for leukocytes such as neutrophils that exhibit a strong exploratory behaviour. A schematic representation illustrating the various measurement methods and the difference they report for identical events is displayed in Figure 8A. Based on end point measurement, we observed that neutrophils migrated further away from the vessel than T-cells in the same time frame, indicating higher migration speed for neutrophils. Using live imaging we found that the effect is even larger than expected due to their exploratory behaviour through the matrix. Live imaging also allowed us to discriminate between migration behaviour of the neutrophil away from but also back towards the vessel. This type of migration behaviour would otherwise not be observed.

**Figure 8.**
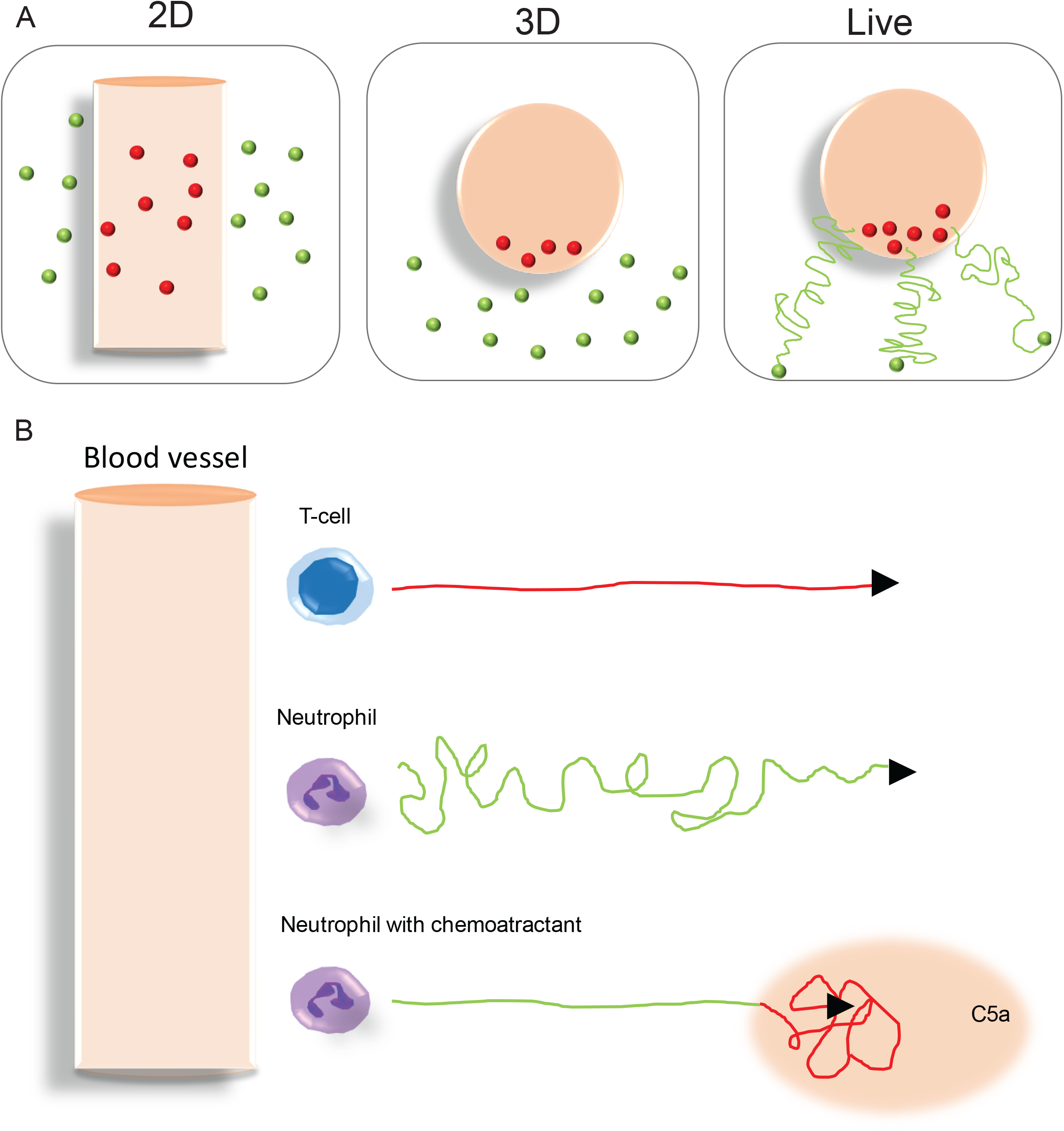
Different imaging methods to study different aspects of leukocyte (trans)migration. (A) Schematic representation of the similar experiment imaged in 2D, 3D or live 3D and with leukocytes perceived to be outside of the vessel indicated as green dots. 2D imaging underestimates the number of extravasated leukocytes because cells underneath the vessel are indistinguishable from cells inside the BVOAC. 3D live imaging allows the tracking of cells over time and therefore analysis of migration dynamics which is not possible based on 3D endpoint imaging. (B) Migration dynamics of different leukocytes in a 3D matrix. T-cells migrate slower and more linear, while neutrophils travel faster while wandering around. When adding a chemoattractant to neutrophils, they initially migrate at a similar speed compared to control, but much more linear. Over time however they lose speed and directionality. Red = slower migration, green = faster migration.

T-lymphocyte migration is characterized by the stop and go manner they display to survey the environment, which makes speed measurements challenging (Dupré *et al.*, 2015; Jerison and Quake, 2020). Sadjadi and colleagues investigated the migration speed of T-cells in different collagen concentrations (Sadjadi *et al.*, 2020). They tested 2, 4 and 5 mg/mL. In our system, we used approximately 2,5 mg/mL and measured a speed of 6 μm/min, comparable with the speed observed by Sadjadi et al. in 2 mg/mL collagen. This is supported by the study of Niggemann et al (1997), who found a migration speed of 7 μm/min in a collagen concentration of 1.67 mg/mL (Niggemann *et al.*, 1997). *In vivo*, higher migration speeds are observed (10-15 μm/min) but this depends on activation status and differs between organs (Miller *et al.*, 2002). Neutrophil migration speed measured in our system is 12 μm/min. A recent study by Wolf et al. measured speed of neutrophils in 3.3 and 8 mg/mL collagen gels and observed migration speeds of 5-10 μm/min respectively (Wolf *et al.*, 2018). Considering that the collagen concentration in our study is slightly lower, the speeds found in our experiments support the speeds mentioned in literature for both T-cells and neutrophils.

The hydrogel-based BVOAC system allows us to study the full TEM process in time in great detail, not hampered by the limitations of classical TEM models. However, there is an important parameter of TEM lacking in this system, which is flow. The importance of flow for endothelial cell function has been shown extensively and is known to affect TEM (Kitayama *et al.*, 2000; Conway *et al.*, 2017; Polacheck *et al.*, 2017). The addition of flow to our BVOAC model would be the next important step, allowing us to investigate the full TEM multistep process from rolling and adhesion, to transmigration and penetration of the collagen matrix in one assay.

Using the BVOAC, we found that neutrophils migrate twice as fast compared to T-cells, and both cell types showed increased migration distance in 3D when first crossing an inflamed endothelial lining. This finding highlights the benefits and opportunities of a BVOAC model over 2D models. Moreover, neutrophils migrate in a random fashion whereas T-cells have a much more directed migration pattern. Under a chemotactic gradient the neutrophil migration adopts more T-cell like directionality for a limited amount of time, before reverting to their exploratory migration pattern. Schematic representations of the various migration dynamics are shown in Figure 8B. Because of the real-time imaging possibility, we found that neutrophils can also migrate back to the vessel once they have entered the matrix. Others have reported on this phenomenon, called reverse transmigration, as well, using *in vivo* animal models (Woodfin *et al.*, 2011; Colom *et al.*, 2015). The BVOAC model now opens new opportunities to study reverse transmigration at the molecular level. This remarkable migration behaviour can also be found when neutrophils do not sense the chemotactic gradient anymore, indicating that the endothelial vessel wall, might in fact attract neutrophils back once the final destination in the tissue has been reached and gradients are cleared. This type of migration is typical for resolving inflammation and was studied using *in vivo* models (Peiseler and Kubes, 2018; Bogoslowski *et al.*, 2020). In this situation the BVOAC model may offer new opportunities to study this migration behaviour as well.

In conclusion, TEM experiments with the BVOAC platform enable the exploration of many parameters previously unobtainable. The level of imaging and analysis determines which aspects of the leukocyte extravasation can be observed. Our comparison between 2D, 3D and live 3D imaging of leukocyte TEM dynamics demonstrates that it is imperative to image and analyse TEM experiments in 3D over time for an accurate interpretation of the events in this system. This hydrogel-based BVOAC model now facilitates studying the full process of human leukocyte extravasation under inflammatory conditions *in vitro* and thereby identify the underlying cellular characteristics and molecular requirements.

## Material and methods

### Cell culture and seeding

Pooled Human Umbilical Vein Endothelial Cells (HUVEC, Lonza P1052, #C2519A) were cultured at 37°C with 5% CO_2_ on fibronectin-coated culture flasks in Endothelial Cell Growth Medium 2 (EGM-2, Promocell #C22011) supplemented with supplement Mix (Promocell, #C39216). HUVECs were passaged at 60-70% confluency and used for experiments between passage 3-7. Pulmonary (HPaMVC, Pelobiotech, # PB-CH-147-4011) and lung (HPuMVC, Lonza, #CC-2527) microvascular cells were cultured at 37°C with 5% CO_2_ on fibronectin-coated culture flasks in SupplementPack Endothelial Cell GM2 containing 0.01 mL/mL L-Glutamine (Sigma, #G3202), 0.02 mL/mL Fetal calf serum (FCS, C-37320), 5 ng/mL human epidermal growth factor (hEGF, C-30224), 0.2 μg/ml Hydrocortisone (HC, C-31063), 0.5 ng/mL Vascular endothelial growth factor 165 (VEGF, C-3260), 10 ng/mL human basic Fibroblast Growth Factor (hbFGF, C-30321), 20 ng/mL Insulin-like Growth Factor (R3, C-31700), 1 μg/mL Ascorbic Acid (AA, C-31750). The cells were passaged at 60-70% confluency and used for experiments between passage 5-8.

### Collagen gel preparations

All following steps were carried out on ice to halt polymerization of the collagen. 50 μl of 10x PBS (Gibco, #70011-044) was added carefully on top of 250 μl of bovine collagen type-1 (10 mg/mL FibriCol, Advanced BioMatrix, #5133) and mixed. When the solution was sufficiently mixed, 48.6 μl of 0.1 M NaOH was added and the mixture was put on ice for 10 minutes. pH was checked before mixing 1:1 with cell culture medium, bringing the final collagen concentration to 2,5 mg/mL.

### 3D vessel production

The LumeNext devices were obtained from the Beebe Lab. For more information about this system, please contact David Beebe. The hollow chambers were coated with 1% Polyethylenimine (PEI, Polysciences, #23966) and incubated for 10 min at room temperature (RT), sequentially chambers were coated with 0.1% glutaraldehyde (Merck, #104239) and washed 5x with Water for Injection (WFI, Gibco, #A12873-01). Chambers were airdried before adding of collagen. Collagen was prepared according to the protocol above. 5 μL collagen was added to the chambers and allowed to polymerize for 30 min. The chamber is about 2mm^3^ in total. PBS drenched cotton balls were added to the device to prevent drying in of the collagen. When polymerization of collagen was confirmed, rods were removed and medium was added to the lumen. HUVECs were dissociated using Trypsin/EDTA (Sigma, #T4049), neutralized 1:1 with Trypsin Neutralising Solution (TNS, Lonza, #CC-5002), spun down and concentrated to 15*10^6^ cells/mL in EGM-2. 5 μl of cell suspension was added per vessel and the devices were placed in a head-over-head rotator for 2 hours at 1 RPM at 37°C with 5% CO_2_ for seeding. Medium was replaced twice daily and vessels were allowed to mature for 2 days before use.

### Neutrophil and T-cell isolation

Polymorphonuclear neutrophils (PMN) were isolated from whole blood derived from healthy donors who signed an informed consent under the rules and legislation in place within the Netherlands and maintained by the Sanquin Medical Ethical Committee. Heparinised whole blood was diluted (1:1) with 10% (v/v) trisodium citrate diluted in PBS. Diluted whole blood was pipetted carefully on 12.5 mL Percoll (RT) 1.076 g/mL. Tubes were centrifuged (Rotanta 96R) at 450 g, slow start, low brake for 20 min. Plasma and peripheral blood mononuclear cell (PBMC) ring fraction were removed. Erythrocytes were lysed in an ice-cold isotonic lysis buffer (155 mM NH4CL, 10 mM KHCO3, 0.1 mM EDTA, pH7.4 in WFI) for 10 min. Neutrophils were pelleted at 450 g for 5 min at 4°C, resuspended in lysis buffer and incubated for 5 min on ice. Neutrophils were centrifuged again at 450 g for 5 min at 4°C, washed once with PBS, centrifuged again at 450 g for 5 min at 4°C and resuspended in HEPES medium (20 mM HEPES, 132 mM NaCl, 6 mM KCL, 1 mM CaCL2, 1 mM MgSO4, 1.2 mM K2HPO4, 5 mM glucose, Sigma-Aldrich, and 0.4% (w/v) human serum albumin, Sanquin Reagents), pH 7.4) and kept at room temperature for no longer than 4 h until use. This isolation method typically yields >95% purity.

Cytotoxic T lymphocytes (CTL) were isolated from density-gradient isolated PBMC by use of magnetic separation. Whole blood was diluted 1:1 with balanced salt solution at RT and layered onto Ficoll Paque PLUS (GE Healthcare, #GE17-1440-02) followed by centrifugation at 400 g for 30 min. From here on cells and buffers are kept at 4°C. The PBMC ring fraction was harvested and washed three times using isolation buffer (PBS + 0.5% FCS). CTL were isolated negatively using the Miltenyi CD8 T-cell isolation kit (Miltenyi, #130-096-495) with LS columns (Miltenyi, #130-042-401) and a QuadroMACS according to the manufacturer’s instruction. Afterwards CTL were harvested and resuspended in RPMI 1640 (Thermo Fisher, #61870010) containing 10% FBS and kept at 37°C and 5% CO2 overnight until use. T-cell purity is typically >95-98% CD8^+^ T cells.

### Immunofluorescent staining

The vessel lumen was washed 3 times with PBS ++ (PBS with 1mM CaCl2, 0.5mM MgCl2) and fixed with 4% paraformaldehyde (PFA, Merck, #30525-89-4) at 37°C for 15 min. Afterwards vessels were incubated with directly conjugated antibodies in PBST (PBS with 0.2% BSA and 0.1% Tween) O/N at 4°C. After staining vessels were washed 3 times with PBS and kept at 4°C awaiting imaging. The following antibodies and stains were used: Mouse anti-hVE-cadherin/CD144 AF647 conjugated ([55-7H1], BD, #561567), Phalloidin AF488 conjugated (Molecular Probes, #A12379), Hoechst 33342 (Molecular probes, #H-1399), Mouse anti-hICAM-1 AF546 conjugated (15.2, Santa Cruz, #sc-107AF546), anti-Collagen IV (Abcam, #ab6586).

### FITC-dextran leakage assay

6 μL of 70 kDa FITC-dextran in EGM-2 (5 mg/mL, Merck, #46945) was added to each vessel. Thrombin (1U/mL, Merck, #T1063) was added to the dextran and injected at the same time. The device was placed under a widefield microscope and every 10 s an image was captured. Images were analysed in Fiji (ImageJ, version 1.52). Fluorescent intensity was measured over the cross-section of the vessel at 6 timepoints throughout the movie.

### TEM assay

Endothelial cells were stained with CellTracker™ Green CMFDA Dye (1 μM, Molecular Probes, #C7025) according to manufactures protocol, before seeding. TNF-α (10 ng/mL, Peprotech, #300-01A) was added O/N inside the vessel. Leukocytes were stained with Vybrant™ DiD Cell-Labelling Solution (1 μM, Molecular Probes) for 15 min at 37°C, pelleted at 450 g for 5 minutes, washed, pelleted again and resuspended in medium. 2 μl of 16*10^6^ neutrophils or T-cells per mL were added per vessel. For live imaging migration was imaged for 60 min and for endpoint conditions neutrophils were allowed to migrate for 2.5 hours at 37°C. Vessels were flushed with PBS++ and fixed for 15 min with 4% PFA. Vessels were stored in PBS at 4°C until imaging.

### Imaging and analysis

Brightfield images were acquired on an Axio Observer Microscope (ZEISS) using Zen 2 blue edition using a 10x air objective (ZEISS, #420341-9911, Plan-Neofluor 10x/0.3 Ph1). Fluorescent staining of the vessels was imaged using a SP8 confocal microscope (Leica) with a 25x long working distance water objective (Leica, #15506375, HC FLUOTAR L 25x/0.95). Control and inflamed vessels were imaged using identical settings. TEM experiments (both live and fixed) were imaged on an LSM980 Airyscan2 (ZEISS) with a 10x air objective (ZEISS, #420640-9900-000, Objective Plan-Apochromat 10x/0.45). Laser power was kept to a minimum to limit phototoxicity during live imaging. The transmigration assays and the 3D rendering/orthogonal views were analysed with IMARIS Bitplane software (Version 9.5/9.6) while the leakage data, zoom-in on the cell junctions and collagen IV stain were analysed with Fiji (ImageJ, version 1.52).

### 3D analysis in Imaris

To analyse the distance from the neutrophils to the surface of the vessel, the spot function was applied to automaticallydetect fluorescently labelled neutrophils and a surface rendering of the vessel was generated from the XYZ-fluorescent label. The vessel surface was created manually by tracing the outline of the vessel, the software then rendered a cylindrical shape corresponding to the immunofluorescent signal. Possible holes were automatically filled by the software. The spot function was applied automatically with the same parameters for all experiments, but checked manually afterwards to ensure detection was correct. When both the vessel surface and spots were created, data like number of neutrophils and distance to the vessels could be extracted from the IMARIS spot function and further analysed in excel.

### Statistics

Statistics were performed in GraphPad Prism 9 (version 9.0.1). Data was checked for normality via the Anderson-Darling test. If normally distributed a (paired) students’ t-test was used to test for statistical significance. Statistical significance is indicated as follows: *= p<0.05, **= p<0.01, ***= p<0.001, ****= p<0.0001.

## Acknowledgements

This work was supported by LSBR grant # 1649 (A.C.I.v.S.), LSBR grant # 1820 (L.K.), NIH R01AI134749 (D.J.B.), NIH R33CA225281 (D.J.B.), NIH P30CA014520 (D.J.B.) and ZonMW NWO Vici grant #91819632 (J.D.v.B.). D.J.B. holds equity in Bellbrook Labs LLC, Tasso Inc., Salus Discovery LLC, Lynx Biosciences Inc., Stacks to the Future LLC, Turba LLC, Flambeau Diagnostics LLC, and Onexio Biosystems LLC. D.J.B. is also a consultant for Abbott Laboratories.

## Author contributions

A.C.I.v.S., L.K. and R.S. performed experiments. M.B., A.C.I.v.S. and L.K. analysed data. M.J.B. advised and provided essential chips. A.C.I.v.S., L.K., J.D.v.B. and M.A.N. wrote manuscript. J.D.v.B. and M.A.N. supervised the study.

