## Supplemental figure for "Transendothelial migration induces differential migration dynamics of leukocytes in tissue matrix"

**Supplemental Figure 1. Neutrophil transmigration during continuous turning.** Representative top view (A) and side view (B) image of neutrophil transmigration in TNF-α treated BVOAC that was placed in a head-over-head during the 2 hour transmigration period. Scalebar = 150 µm.

**Supplemental video 1. Neutrophil transmigration through the endothelial wall of the BVOAC in high resolution over time.** The video is a blended projection of 3D imaging over time of neutrophils extravasating from the lumen of the BVOAC, through the vessel wall and into the collagen matrix surrounding the vessel. The video rotates the scene to give an overview of the red neutrophils in the lumen of the vessel on top of the green endothelial cell layer. Green blobs inside the lumen are detached and rounded up HUVEC. At 10 seconds into the video the experiment starts playing and 2 red neutrophils start poking through the vessel wall after which the time is paused and the scene is rotated to show the outside of the endothelial layer with the neutrophils poking through. The scene then changes back to inside the lumen and resumes playing at 18 seconds showing the neutrophils both crossing the barrier layer until finally they have completely crossed the endothelium and are seen migrating at the basolateral side of the endothelium. Time elapsed in the experiment is shown in the bottom right corner of the video and the adaptive scale bar is shown in the bottom left of the video

**Supplemental video 2. Data of experiment shown in supplemental video 1 with rendered surfaces to further reveal details of the TEM event.** The video shows the raw data in a MIP projection and rotates the scene 360 degrees first before zooming in on the location where 2 neutrophils sequentially migrate at the same junction. The rendered volumes are projected on top of the MIP projection of the data at 9 seconds into the video, to clarify outlines of the EC and the 2 neutrophils focussed in this rendering. The video then plays the experiment over time until 17 seconds where the neutrophil is poking through the endothelial layer. The video stops the time and rotates to show the leading edge of the neutrophil in the matrix outside of the endothelial barrier. The video afterwards resumes displaying the experiment over time. Time elapsed in the experiment is shown in the bottom right corner of the video and the adaptive scale bar is shown in the bottom left of the video.

**Supplemental video 3. Neutrophils migrating out of the BVOAC lumen into the collagen matrix imaged in 3D over one hour displayed from a side view.** This experiment shows the raw data of red labelled neutrophils migrating out of the CellTracker green stained HUVEC vessel rendered in transparent green. HUVEC were treated overnight with TNF-α to induce inflammation. To capture as many neutrophils as possible the bottom part of the vessel and a large part of the collagen matrix were imaged. To illustrate the path travelled by a single neutrophil, the track is shown in a colour coded line which represents the time passed in the experiment. Scale bar for size is shown in the left bottom corner of the video and time including colour legend is shown in the bottom right of the video.

**Supplemental video 4. T-cells migrating out of the BVOAC lumen into the collagen matrix imaged in 3D over one hour displayed from a side view.** This experiment shows the raw data of red labelled T-cells migrating out of the CellTracker green stained HUVEC vessel rendered in transparent green. HUVEC were treated overnight with TNF-α to induce inflammation. To capture as many events as possible the bottom part of the vessel and a large part of the collagen matrix were imaged. To illustrate the path travelled by a single T-cell the track is shown in a colour coded line which represents the time passed in the experiment. Scale bar for size is shown in the left bottom corner of the video and time including colour legend is shown in the bottom right of the video.

**Supplemental video 5. Neutrophils migrating out of the BVOAC lumen into the collagen matrix under the influence of c5a imaged in 3D over one hour.** This experiment shows the raw data of red labelled T-cells migrating out of the CellTracker green stained HUVEC vessel rendered in transparent green. The experiment is displayed from a top view with the vessel directed horizontally and the matrix inlets located to the top and bottom of this view. A C5a gradient was created in this BVOAC by adding c5a to one of the two matrix inlets of the device. This inlet is located towards the bottom side of this video. To illustrate the path travelled by a single neutrophil the track is shown in a colour coded line which represents the time passed in the experiment. The vast majority of neutrophils migrates towards the C5a gradient for the first half of the video after which this directionality is lost and the neutrophils assume random directionality. This is also reflected in the highlighted track. Scale bar for size is shown in the left bottom corner of the video and time including colour legend is shown in the bottom right of the video.

Supplemental Figure 1.

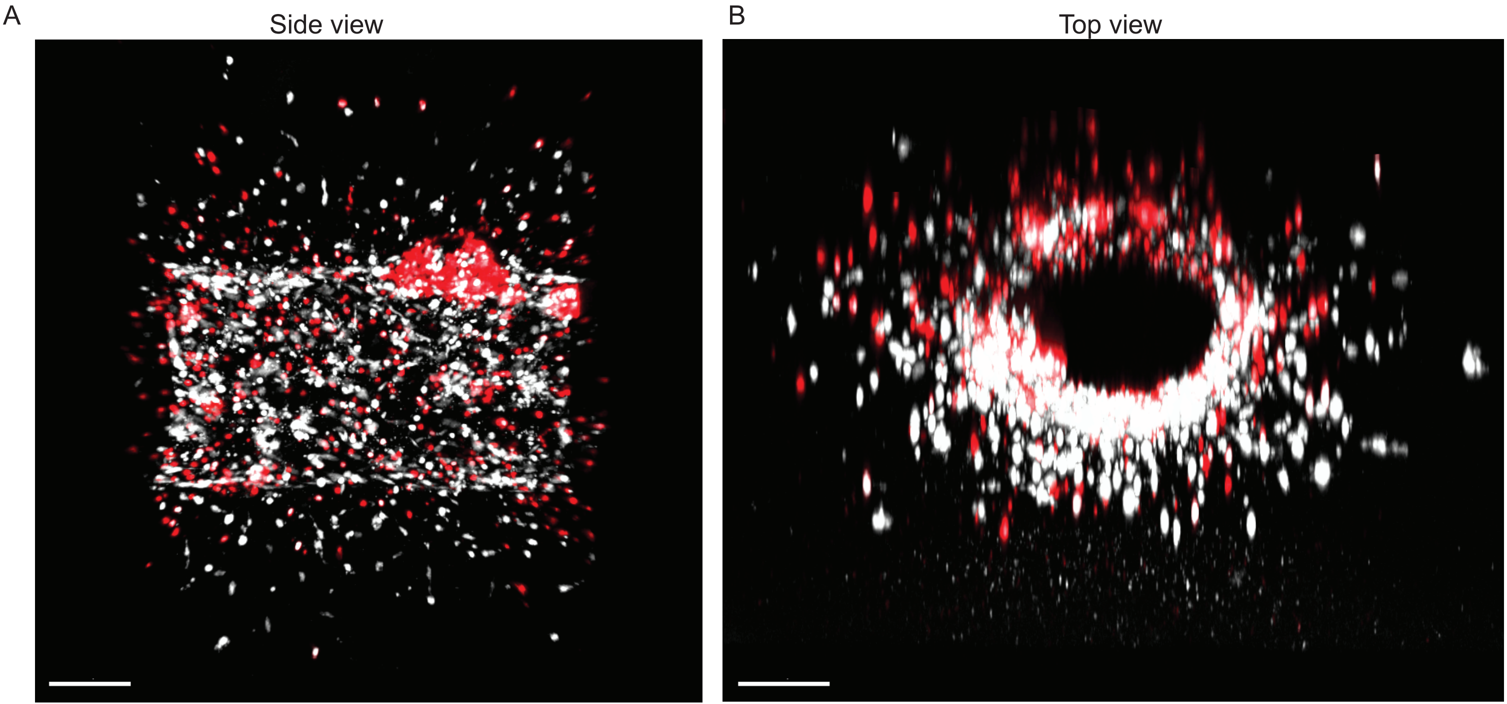
